# Reproducing Transformation of Indolent B-cell Lymphoma by T-cell Immunosuppression of L.CD40 Mice

**DOI:** 10.1101/477273

**Authors:** Christelle Vincent-Fabert, Alexis Saintamand, Amandine David, Mehdi Alizadeh, François Boyer, Nicolas Arnaud, Ursula Zimber-Strobl, Jean Feuillard, Nathalie Faumont

**Author notes:** **Corresponding Author:** Nathalie Faumont; CNRS-UMR 7276 INSERM U1262 CRIBL "Contrôle de la Réponse Immune B et Lymphoproliférations", CBRS “Centre de Biologie et de Recherche en Santé”, Dupuytren Hospital University Center, University of Limoges, Hematology Laboratory of Dupuytren CHU, 2 rue du Docteur Marcland, 87025 Limoges, France.; and.

## Abstract

Transformation of an indolent B-cell lymphoma is associated with a more aggressive clinical course and poor survival. The role of immune surveillance in the transformation of a B-cell indolent lymphoma towards a more aggressive form is poorly documented. To experimentally address this question, we used the L.CD40 mouse model, which is characterized by B-cell specific continuous CD40 signaling, responsible for spleen indolent clonal or oligoclonal B-cell lymphoma after one year in 60% cases. Immunosuppression was obtained either by T/NK cell depletion or by treatment with the T-cell immunosuppressive drug cyclosporin A. Immunosuppressed L.CD40 mice had larger splenomegaly with increased numbers of B-cells in both spleen and peripheral blood. High-throughput sequencing of immunoglobulin variable segments revealed that clonal expansion was increased in immunosuppressed L.CD40 mice. Tumor B cells of immunosuppressed mice were larger with an immunoblastic aspect, both on blood smears and spleen tissue sections, with increased proliferation rate and increased numbers of activated B-cells. Collectively, these features suggest that immune suppression induced a shift from indolent lymphomas into aggressive ones. Thus, as a preclinical model, immunosuppressed L.CD40 mice reproduce aggressive transformation of an indolent B-cell tumor and highlight the role of the immune surveillance in its clinical course, opening new perspective for immune restoration therapies.

**Summary statement:** Highlighting the role of immune surveillance, transformation of indolent B-cell lymphoma into an aggressive malignancy is experimentally reproduced after T-cell immune suppression in the L.CD40 preclinical mouse model.

## Introduction

Indolent B-cell lymphomas are a group of incurable cancers encompassing various entities such as chronic lymphocytic leukemia (CLL), follicular lymphoma (FL) or marginal zone lymphoma (MZL). They occur predominantly in elderly subjects and are extremely rare in patients younger than 40 years old. Various alterations of the immune system are associated with aging including a shift from a naive to a memory T-cell phenotype, decreased T-cell responses to *in vitro* stimulation, and oligoclonal expansion in the T- and B-cell repertoire (Linterman, 2014; Nikolich-Zugich, 2008; Sarkozy et al., 2015). Aging has been shown to influence the expression of co-stimulatory molecules such as ICOS and CTLA-4 expression on T-cells (Canaday et al., 2013) and is also associated with increased frequency of regulatory T-cells (Treg) in both mice and humans. As Tregs control the intensity of T-cell responses, they could contribute to age-related immune dysfunction (Raynor et al., 2012). Consequences of immune decline in elderly people include lower vaccination efficacy, decreased resistance to infections, increased inflammation and autoimmune activation, decreased immune surveillance and increased onset of malignancies (Ponnappan and Ponnappan, 2011; Pinti et al., 2016).

Despite these immunological deficits in elderly individuals, immune surveillance still remains, and the indolent tumor B-cells are very likely to subvert their microenvironment with a prolonged escape phase. For example, FL B-cells may secrete CCL22, which recruits T regulatory cells (Treg). Furthermore CD4 positive T-cells from FL follicles exhibit a profile of exhausted effector T-cells with a high proportion of PD1 and/or TIM3 positive cells (Amé-Thomas and Tarte, 2014). Immune suppression and poor antitumor immune responses are among hallmarks of CLL (García-Muñoz et al., 2012). CD5, the main surface immunophenotypic marker of CLL, can lead to IL10 secretion (Gary-Gouy et al., 2002), which acts not only as an autocrine growth factor for leukemic B-cells, but also as an immunosuppressive cytokine inhibiting T-cells and antigen presenting cells (Ramsay et al., 2008). Standardized incidence ratios of splenic marginal zone lymphomas is markedly increased in young adults after solid organ transplantation (Clarke et al., 2013), a clear indication that immune surveillance may prevent emergence of these lymphomas. Consistently, the presence of PD-1 positive exhausted T-cells has been reported in marginal zone lymphomas (Xu-Monette et al., 2017). All indolent lymphomas may evolve towards aggressive transformation with an increased proliferation index and decreased tumor doubling time. Such transformation, called Richter’s syndrome in CLL, is associated with a more aggressive clinical course and poor survival. Indeed, progression of indolent B-cell lymphomas is likely to be associated with escape from immune surveillance(Nicholas et al., 2016). It is of note, therapies targeting the PD-1/PD-L1 axis seem to be effective only on CLL with Richter’s transformation (Ding et al., 2017). However, in fact, the role of the immune system is poorly known in such transformation process.

Our aim was to experimentally address the question of the role the immune suppression in transformation of an indolent B-cell lymphoma. Among the very few mouse models of indolent lymphoma is the one published by Hömig-Hölzel *et al* (Hömig-Hölzel et al., 2008). In this model, the transgene, preceded by a loxP flanked stop-cassette in the *rosa26* genomic locus, encodes for a chimeric protein composed of the transmembrane domain of latent membrane protein 1 (LMP1) of EBV and the intracellular signaling domain of CD40, *i.e.* L.CD40 mice. This chimeric protein is responsible for a continuous CD40 signal, which results in continuous NF-_Ƙ_B activation (Gires et al., 1997). When crossed with mice expressing Cre recombinase under control of the CD19 promoter (CD19-Cre mice), L.CD40 mice first exhibit an expansion of the marginal B-cell compartment in the spleen and then develop an indolent lymphoma of the spleen after one year in 60% of cases (Hömig-Hölzel et al., 2008). Construction of the L.CD40 model is closed from the one of Zhang *et al* in which the transgene was the latent membrane protein 1, the main oncogenic protein of Epstein Barr Virus (EBV), which acts by rerouting CD40 signaling pathways (Zhang et al., 2012). In this model, LMP1 expressing B-cells were almost completely eliminated by the host immune system, and only deep T and NK cell depletion allows rapid emergence of highly aggressive B-cell lymphomas.

Here, we found that immune suppression either by T-cell depletion or by treatment with the T-cell immunosuppressive drug of L.CD40 mice induced a shift from an indolent to an aggressive lymphoproliferative disorder. These results suggest that, according to the immune status, the L.CD40 mouse model could be a preclinical model not only of indolent B-cell lymphoma but also of their transformed counterpart and point on the role of immune anti-tumor surveillance in the natural course of these cancers.

## Results

### Increased splenomegaly is due to spleen expansion of B-cells in immunosuppressed L.CD40 mice

Raising the question of the role of immune surveillance in indolent B-cell lymphomagenesis, we tested the effect of immunosuppression on indolent lymphoma developed by L.CD40 mice. Eight months old L.CD40 mice were treated either every 4 days with a cocktail of monoclonal antibodies (mAbs) against T and NK-cells for 3 weeks or daily with CsA for 3 weeks or 3 months. T-cell depletion and CsA treatment had no effect on the size and weight of spleens from CD19-Cre mice (not shown). After three weeks treatment with mAbs against T and NK cells, T-cell depletion was very pronounced in spleen and almost complete in blood (Figure S1A and S1B). After CsA treatment, T-cells strongly decreased in the blood and only mildly in the spleen (Figure S1C). T and NK cell depletion by mAbs was associated with increased spleen enlargement (Figure 1A). This spleen enlargement was related to increase in splenocyte absolute numbers (Figure 1B), that was mainly due to B-cell compartment expansion (Figure 1C). Three weeks immunosuppression with CsA had no significant effect on spleen size (not shown). After 3 months, spleens of control L.CD40 mice were further enlarged due to aging (Figure 1A). However, three months immunosuppression with CsA induced a more pronounced splenomegaly (Figure 1A), with a significant increase in absolute number of spleen cells (Figure 1B), that was also due to increased spleen B-cell content (Figure 1C).

**Figure 1:**
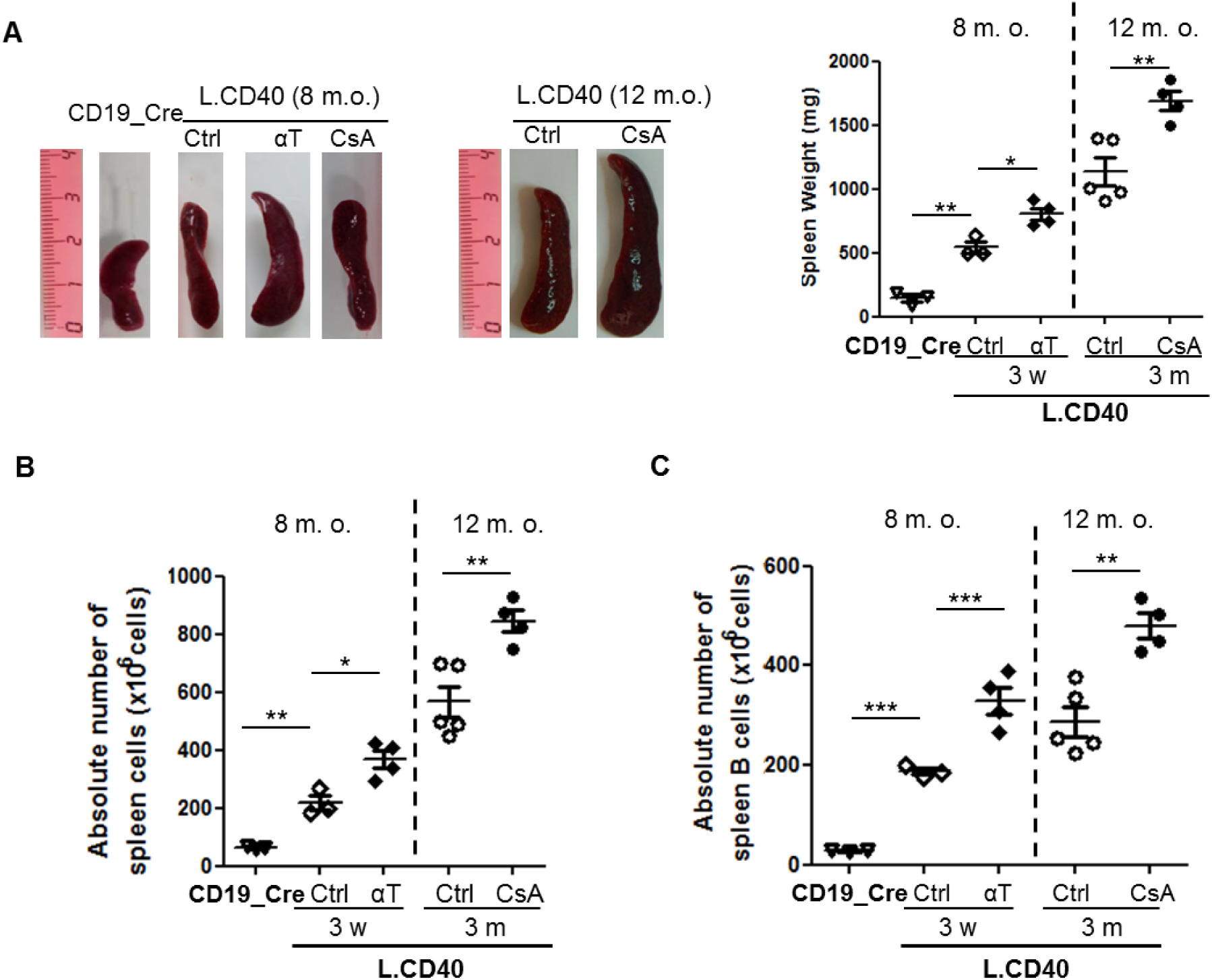
Increased B-cell numbers in immunosuppressed L.CD40 mice. L.CD40 mice were immunosuppressed through intra-peritoneal injection of anti-T cocktail (L.CD40 T) or CsA (L.CD40 CsA). CD19_Cre mice were used as control wild type mice. For each treatment, isotype antibody (for T cell depletion), or placebo (for CsA treatment) were used as controls (Ctrl). (A) Representative spleen size (left panel), and spleen weight (right panel) of L.CD40 mice immunosuppressed by αT (3 weeks; w) or CsA (3 months; m). (B) Absolute numbers of spleen cells in L.CD40 mice after depletion of T cells for 3 weeks (w) or after CsA treatment for 3 months (m). (C) Flow cytometry of B220 spleen B-cell absolute numbers in immunosuppressed L.CD40 mice. Age of mice at the end of treatment is indicated above dot plots (m.o.; month old). Statistical significance was determined by Student’s t-test (*** *p<0.001*; ** *p<0.01*; * *p<0.05*).

### Increased spleen expansion of B-cells in immunosuppressed L.CD40 mice is related to increased B-cell clonal abundance with presence of activated large cells and increased proliferation index

Analysis of B-cell clonality was done by high-throughput sequencing of VDJ regions (Figure 2). With a threshold of 10% clonal frequency (or clonal abundance), no significant clonal expansion was seen in one year old control wild type mice. Even if spleen morphology remained globally unchanged, without splenomegaly and without B-cell expansion, three-month CsA treatment of control mice was associated with presence of spleen B-cell clones above 10% in one out of the three tested cases (33%) cases. Three out of the five (60%) untreated L.CD40 mice exhibited clonal B-cell expansion. Five out of five (100%) three month CsA immunosuppressed L.CD40 mice were clonal. Mean clonal abundance of the dominant clone was 42% and 12% in L.CD40 mice with or without immune suppression respectively (Student T-test, p = 0.03), without any bias in terms of V segment usage (not shown). These results indicate that long term CsA induced immunosuppression did significantly favor expansion of clonal B-cells in L.CD40 mice. Morphologically, spleen B-cell increase in three-month CsA immunosuppressed mice was associated with broad sheets of large cells with oval nuclei, lacy chromatin and paracentral nucleoli (Figure S2A). This cell size enlargement was confirmed by flow cytometry since forward scatter of B-cells was increased in immunosuppressed mice (Figure S2B). Expression of activation markers such as CD80 and CD86 was up-regulated in IS L.CD40 mice (Figure 3A). The fraction of BrdU positive B-cells from immunosuppressed L.CD40 mice was increased when compared to controls (Figure 3B) as well as numbers of Ki67 positive cells in spleens (Figure S2C). Altogether, these results suggest that long-term immune suppression was associated with both increased clonal expansion and aggressive transformation of the indolent L.CD40 B-cell lymphoma.

**Figure 2:**
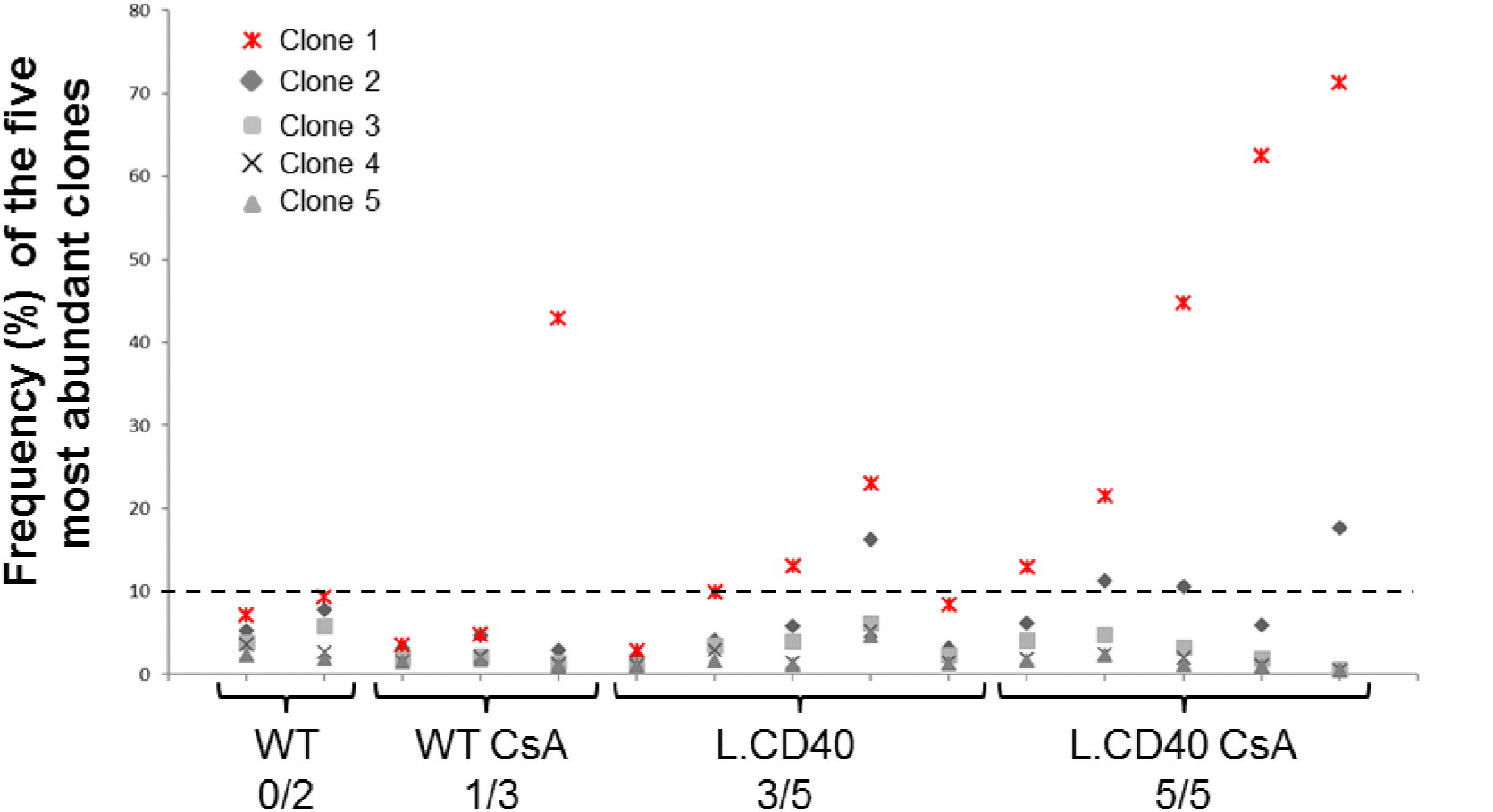
Frequency of the five most abundant clones for wild type (WT) and L.CD40 mice treated or not with CsA. Legends of clones are ordered by decreasing frequency for each mouse. The dominant clone is highlited in red

**Figure 3:**
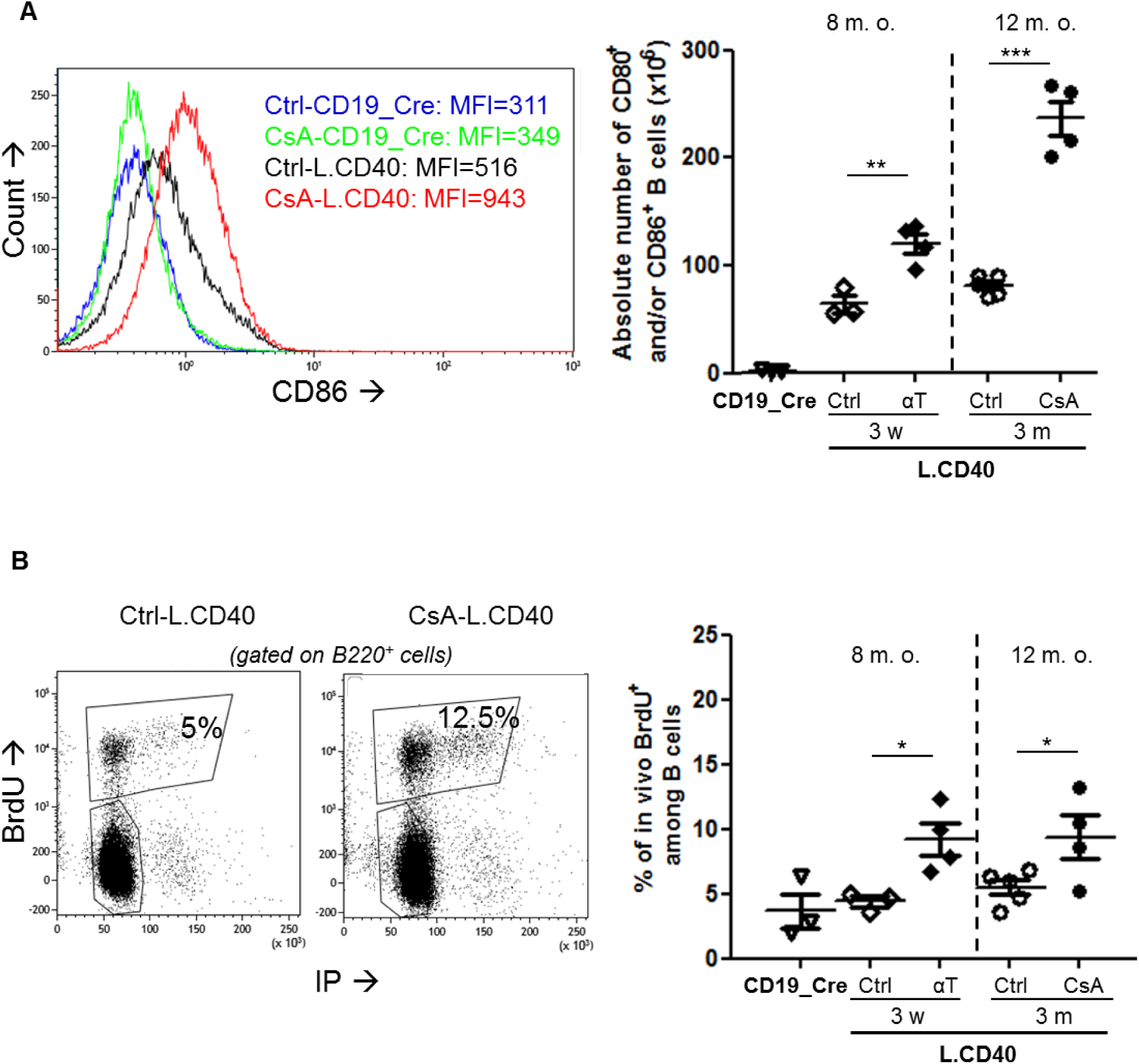
Gain of proliferation and activation in spleen B-cells from L.CD40 immunosuppressed mice. (A) Representative overlay graph for the cell surface expression of marker CD86 on spleen B-lymphocytes (left panel); means and standards deviation of absolute numbers of B220 spleen B-cells expressing activation markers CD80 and/or CD86 (right panel). (B) Flow cytometry graph of BrdU and DNA content by propidium iodide staining for Ctrl or CsA L.CD40 (left panel); Means and standards deviation of flow cytometry percentages of BrdU positive B-cells after *in vivo* BrdU incorporation (right panel). Age of mice at the end of treatment is indicated above dot plots (m.o.; month old). Statistical significance was determined by Student’s t-test (*** *p<0.001*; ** *p<0.01*; * *p<0.05*).

### Increased amounts of large B-cells in the blood of long term immunosuppressed L.CD40 mice

No significant changes were seen in blood from three weeks T and NK cell depleted mice (not shown). After two months, CsA treatment we observed increased leukocytes in L.CD40 mice (Figure 4A). As assessed by flow cytometry, the B-cell compartment of CsA treated L.CD40 mice was increased, contrasting with the decrease in granulocytes and T-cells (Figure 4B). We also noticed the emergence of large B-cells on the forward scatter (FCS) monoparametric histograms (Figure 4C left panel). This cell size increase was confirmed on blood smears after May-Grunwald Giemsa (MGG) staining (Figure 4D right panel). Lymphocytes from control L.CD40 mice remained small with little cytoplasm, round nuclei and dense chromatin while those from CsA treated L.CD40 mice were often large with abundant basophilic cytoplasm, and prominent nucleoli.

**Figure 4:**
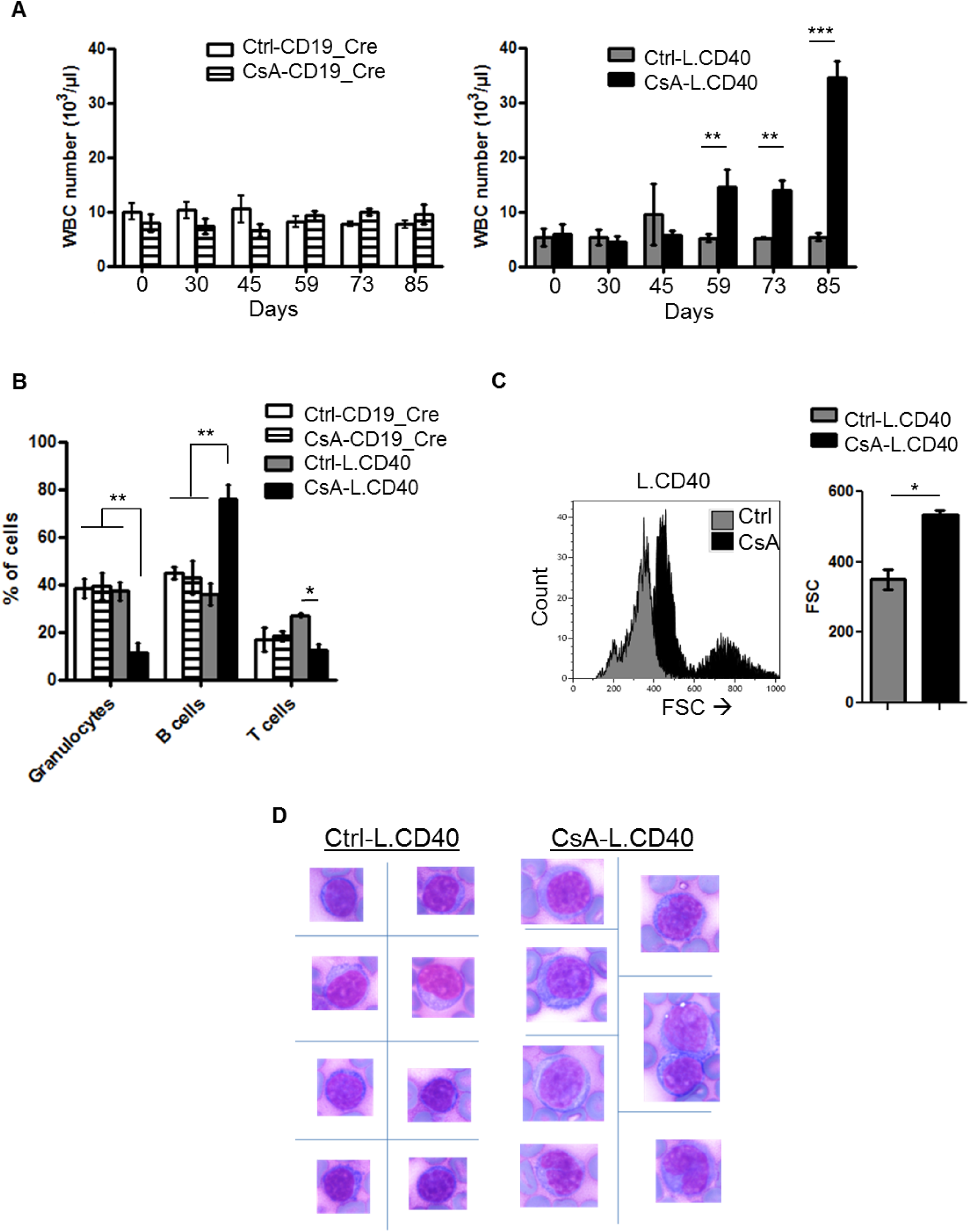
B-cell expansion into blood in long term CsA immunosuppressed L.CD40 mice. (A) Number of circulating white blood cells in immunosuppressed (CsA) or untreated (Ctrl) CD19_Cre and L.CD40 mice. (B) Flow cytometry percentages of circulating granulocytes (Gr-1 positive cells), B-cells (B220 positive cells) and T-cells (CD3 positive cells). Each mouse type and treatment condition is indicated at the top right of the graph. (C) Flow cytometry estimation of circulating B-cell size from Ctrl and CsA-L.CD40 mice. Left panel, overlay of FSC monoparametric histograms gated on B220 B-cells. Right panel, means and standard deviations of FSC for all studied mice. (D) Representative lymphocyte morphology after May Grünwald Giemsa staining of blood smears from Ctrl and CsA-L.CD40 mice (magnification X 1000). For all experiments, at least 4 mice were studied for each condition. Statistical significance was determined by Student’s t-test (*** *p<0.001*; ** *p<0.01*; * *p<0.05*).

## Discussion

Here, we experimentally raise the question of role of the immune suppression in the transformation of an indolent B-cell lymphoma using the L.CD40 mouse model. It was seen that immune suppression accelerated clonal B-cell expansion and increased the tumor aggressiveness.

Post-transplant lymphoproliferative disorders is one of the main clinical situation in which therapeutic induced immune suppression (CsA being one of the main immune suppressive drug) is the direct cause of B-cell lymphoma. These B-cell lymphomas are almost always aggressive and are very often associated with EBV. Indeed, the vigorous immune surveillance exerted against EBV-immortalized B-cells in immunocompetent hosts constrains the EBV-immortalized B-cell to adopt a silenced non-proliferating phenotype to evade the immune response. Such role of the immune surveillance has been experimentally demonstrated by Zhang *et al* who show, in a genetically modified mouse model, that the immune system is able to eliminate B-cells that have been endogenously activated by LMP1, the main oncoprotein of EBV (Zhang et al., 2012). In both cases of EBV infection or LMP1 mouse model, the immunocompetent host is able to completely abolish emergence of clonal B-cell lymphoma, and only immune suppression reveals the transformation potential of EBV and its oncoprotein LMP1. The L.CD40 model is different since B-cells bearing CD40 continuous signaling slowly accumulate with late clonal emergence. Moreover, after immune suppression, emergence kinetics of aggressive B-cell clones is much faster for the LMP1 mouse model than for the L.CD40 model, taking few weeks for the former(Zhang et al., 2012) and few month for the latter (our results). In that view B-cells bearing CD40 continuous signaling are very likely to be able to escape from the host immune system and to progressively accumulate. CD40 and LMP1 signaling pathways are not exactly similar (Graham et al., 2010). For example, CD40 activation of the alternate NF-κB pathway is more intense than the one of LMP1 that mainly activates the classical pathway (Chanut et al., 2014).

A pivotal event in the natural history of almost all indolent B-cell lymphomas is transformation into a more aggressive malignancy, most frequently resembling DLBCLs. At 5 years, transformation frequency may be below 5%, as for marginal zone lymphomas and over 20% for FCLs and CLL (Montoto and Fitzgibbon, 2011). Among genetic events associated with transformation are those promoting c-Myc activation or TP53 inactivation(Lossos and Gascoyne, 2011). With the background of a primary genetic event affecting the common precursor tumor cells such as translocation of BCL2 in FCLs, del(13q) or del(12q) in CLL and MALT1 or BCL10 in MALT marginal zone lymphomas, acquisition of genetic events promoting proliferation and/or resistance to cell death would favor Darwinian selection of more aggressive subclones. Such a transformation arises either from the dominant clone as in most cases of CLL (Fabbri et al., 2013; Mao et al., 2007) or from clonally related common precursor tumor cell as in FCLs (Bouska et al., 2017; Fabbri et al., 2013; Pasqualucci et al., 2014). Transformation of FL is associated with genetic events favoring immune escape (Bouska et al., 2017; Fabbri et al., 2013; Pasqualucci et al., 2014). In solid cancers such as melanomas or lung carcinomas, there is a clear relationship between the mutation burden and a positive response to immunotherapies directed against the PD-L1/PD-1 axis, most likely due to the high number of tumor specific neo-antigens (Simone et al., 2015). Acquisition of additional genetic events in indolent lymphomas would be associated with increased immunogenicity of the subclonal B-cells. As noted above, anti PD-1 immune therapies are effective only in CLL with Richter’s transformation. In this context, it can be put-forward that transformation of an indolent B-cell lymphoma could be due either to aggravation of patient immune deficiency or to genetic acquisition of a series of tools able to neutralize the anti-tumor immune response by the transformed B-cell while increasing its proliferation index.

L.CD40 indolent B-cell lymphomas are characterized by increase in splenomegaly and considerable expansion of splenic B-cells with late acquisition of oligo or monoclonality. The proliferative index remains consistently low or moderate, with no evidence for transformation into high-grade lymphoma. Immunosuppression of L.CD40 mice led to increased aggressiveness of the lymphoproliferative B-cell disorder. Signs of aggressiveness included increased splenomegaly, size and proliferative index of cells, expression of activation markers and blood passage of B-cells. This transformation was associated with increased expansion of B-cell clones. It seems likely that immune suppression was be associated with a morphological, immunophenotypic and proliferative shift toward tumor aggressiveness from preexisting clones. This suggests that induced immunosuppression removed a control exerted by the immune system on B-cell expansion in L.CD40 mice and reproduce the main features of indolent B-cell lymphoma transformation. Thus, in the view of therapies allowing immune restoration, the L.CD40 mouse model opens interesting perspectives not only as preclinical model of indolent B-cell lymphoma but also as a model of aggressive transformation of a B-cell indolent clone.

## Material and Methods

### Mouse models and treatments

L.CD40 mice have been already described (Hömig-Hölzel et al., 2008). Animals were housed at 21–23°C with a 12-h light/dark cycle. All procedures were conducted under an approved protocol according to European guidelines for animal experimentation (French national authorization number: 87-022 and French ethics committee registration number “CREEAL”: 09-07-2012). For antibody-mediated T-cell depletion, L.CD40 mice were injected intraperitoneally every 4 days for three weeks with a mix of anti-CD4 (YTS 191.1.2), anti-CD8 (YTS 169.4.2.1), and anti-Thy-1 (YTS 154.7.7.10) antibodies (200 µg each) in In VivoPure Dilution Buffer, (Bio X cell; US). For CsA treatment, wild type and L.CD40 mice were injected intraperitoneally daily for three months with placebo (diluent composed of ethanol and cremophor) or 10 mg/kg CsA (Sandimmun – Novartis; US) diluted in 5% glucose.

### Flow cytometry

Blood from mice (200 μL) was collected intra-orbitally. Spleen and lymph nodes were also collected and immune cells were filtered through a sterile nylon membrane. Cell suspensions were stained at 4°C in FACS Buffer (PBS, 1% FBS, 2 mM EDTA). The following fluorescent-labelled antibodies were used: anti B220-BV421 (clone RA3-6B2, 1/400), anti CD80-APC (clone16-10A1, 1/2500), anti CD86-FITC (clone GL-1, 1/600), anti CD3-PE (clone 17A2, 1/100) from BioLegend (San Diego, CA, USA). Stained cells were analyzed using a BD-Fortessa SORP flow cytometer (BD Bioscience; US). Results were analyzed using Kaluza Flow Cytometry software 1.2 (Beckman Coulter; France).

### Proliferation

For in vivo proliferation, mice were injected intraperitoneally with 2 mg BrdU (Sigma-Aldrich, US), 18 hours before isolating cells. Splenocytes were stained for B220 and phases of cell cycle were analyzed by measuring BrdU and Propidium Iodide (PI)-incorporation, using the FITC-BrdU Flow Kit (BD Pharmingen; US).

### Sequencing of VDJ regions

RNA was extracted from total spleen, and one μg was used for sequencing. Immunoglobulin gene transcripts were amplified by 5’RACE PCR as described (Boice et al., 2016). Illumina sequencing adapter sequences were added by primer extension, and resulting amplicons were sequenced on an Illumina MiSeq sequencing system using MiSeq kit Reagent V3 600 cycles. Repertoire analysis was done using IMGT/HighV-QUEST tool and R software. Briefly, VH, JH and CDR3 segments were identified using HighV-QUEST. Based on these annotations, reads were grouped in clonotypes that share the same VH and JH gene and high CDR3 homology. The relative abundance of each clonotype was then calculated.

## Acknowledgments

We thank Dr J Cook Moreau, UMR CNRS 7276, Limoges, France, for English editing. The group of J Feuillard is supported by grants from the Ligue Nationale contre le Cancer (Equipe Labellisée Ligue), the Institut National contre le Cancer (INCa), the Comité Orientation Recherche Cancer (CORC), the Limousin Region and the Haute Vienne and Corrèze committees of the Ligue Nationale contre le Cancer and by the Lyons Club of Corrèze. U Zimber-Strobl was supported by the German Research Foundation (SFB 1243, TP-A13 and ZI1382/4-1).

## Authorship Contributions

C.V.F. performed and analyzed experiments, and contributed to the writing. N.A. and A.D. helped to perform proliferation and flow cytometry experiments. A.S., M.A, and F.B. analysed the B-cell repertoire. U.Z.S. helped analyze the results and contributed to the writing of the manuscript. J.F. and N.F. designed and directed the study, contributed to the experiments, analyzed the results and wrote the manuscript.

